# Major effect loci for plant size before onset of nitrogen fixation allow accurate prediction of yield in white clover

**DOI:** 10.1101/2021.04.16.440135

**Authors:** Sara Moeskjær, Cathrine Kiel Skovbjerg, Marni Tausen, Rune Wind, Niels Roulund, Luc Janss, Stig U. Andersen

**Affiliations:** Department of Molecular Biology and Genetics, Aarhus University, 8000 Aarhus C, Denmark; Bioinformatics Research Centre, Aarhus University, 8000 Aarhus C, Denmark; DLF, 4660 Store Heddinge, Denmark; Center for Quantitative Genetics and Genomics, Aarhus University, 8000 Aarhus C, Denmark

**Keywords:** *Trifolium repens*, nitrogen fixation, yield, GWAS, genomic prediction, genomic selection, genetics, sustainable agriculture, symbiosis

## Abstract

White clover is an agriculturally important forage legume grown throughout temperate regions as a mixed clover-grass crop. It is typically cultivated with low nitrogen input, making yield dependent on nitrogen fixation by rhizobia in root nodules. Here, we investigate the effects of clover and rhizobium genetic variation by monitoring plant growth and quantifying dry matter yield of 704 combinations of 145 clover and 169 rhizobium genotypes. We find no significant effect of rhizobium variation. In contrast, we can predict yield based on a few white clover markers strongly associated with plant size prior to nitrogen fixation, and the prediction accuracy for polycross offspring yield is remarkably high. Several of the markers are located near a homolog of *Arabidopsis thaliana GIGANTUS 1*, which regulates growth rate and biomass accumulation. Our work provides fundamental insight into the genetics of white clover yield and identifies specific candidate genes as breeding targets.

## Introduction

White clover (*Trifolium repens* L.) is an important forage crop in temperate climates. It improves forage quality by increasing protein content, digestibility and palatability in perennial grass pastures and allows reduced nitrogen fertilizer input due to symbiotic nitrogen fixation with rhizobia (Archer, 1973; Ruz-Jerez et al., 1991; Thomson et al., 1985). It is a relatively young, outcrossing species, which originated during the most recent glaciation around 20,000 years ago by hybridisation of two diploid species, *T. occidentale* and *T. pallescens* (**Figure 1A**) (Griffiths et al., 2019).

**Figure 1.**
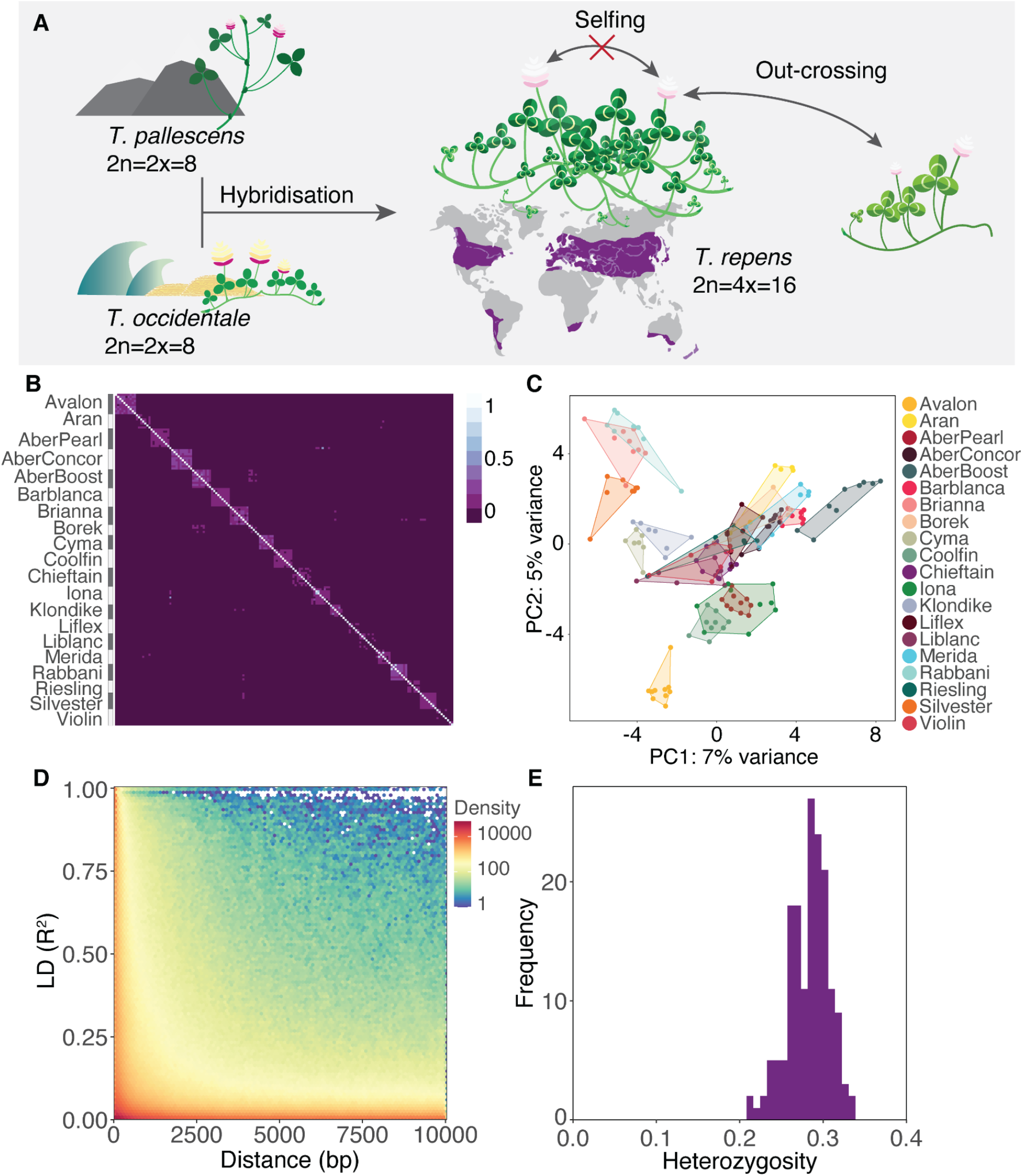
Characterization of the white clover population. **A:** White clover origin, range, and out-crossing mating habit. **B:** Heat map of genomic relationship matrix (GRM) for the 148 clover genotypes. **C:** Population structure of the 148 clover genotypes by the first two principal components of the GRM. **D:** LD (*R^2^*) for the RNAseq SNP dataset. **E:** Heterozygosity of individuals.

In grass-clover pastures, three main components and the interaction between them determine yield: clover, grass and rhizobia. Under low nitrogen input, yield can improve many-fold if legumes are inoculated with the appropriate symbiont, supporting sustainable agricultural systems to feed livestock (Caradus et al., 1995). Since the 1970s, a large number of studies have investigated the interactions between white clover and rhizobium. Many examples of successful inoculation have been reported in locations where the natural occurrence of *Rhizobium leguminosarum* sv. *trifolii* white clover symbionts is low (Irisarri et al., 2019; Lowther & Kerr, 2011; M. M. Svenning et al., 2001; Young & Mytton, 1983), and a number of examples of white clover-rhizobium interactions that affect yield exist (Mytton, 1975; Mette M. Svenning et al., 1991; Young & Mytton, 1983), although other studies reported only small effects or no such interactions (Crush, 1995).

A major objective of white clover breeding is to improve the biomass yield and thus reduce the land use required for supporting meat and dairy production (Hayes et al., 2013), but the genetic gain for dry matter yield in clover has increased at a moderate to low rate in the last 90 years (Hoyos-Villegas et al., 2019). Currently, white clover is primarily bred using phenotypic selection, which might be confounded by large effects of phenotypic plasticity, limiting accurate estimates of breeding values, and requires relatively long generation intervals as the breeder has to wait for traits to become observable (Hayes et al., 2013; Hoyos-Villegas et al., 2019). Although several studies have reported successful prediction of complex traits when applying genomic selection (GS) to important crops (Voss-Fels et al., 2019), the application of genetic markers in breeding of white clover has been very limited, probably because of its complex genetic nature (Faville et al., 2012). However, with the emergence of a high quality reference genome with extensive gene annotation (Griffiths et al., 2019), applying genomics in the breeding practices of white clover has become more attractive, especially considering that GS has shown promising results for biomass yield in alfalfa, an autotetraploid forage legume (Paolo Annicchiarico et al., 2015).

Identification of quantitative trait loci (QTLs) can help accelerate the yield improvement of future cultivars. During the last decades efforts have been made to identify QTLs associated with yield in important crops (Bernardo, 2008). Examples of these studies include the identification of QTLs that explain between 5% and 45% of the variation in plant biomass yield in rye, sweet potato, rice and alfalfa using family-based linkage studies or genome-wide association studies (GWAS) (Matsubara et al., 2016; Miedaner et al., 2018; Sakiroglu & Brummer, 2017; Zhao et al., 2013).

Most traits associated with agronomically important traits such as yield are controlled by many loci each contributing small effects (Bernardo, 2008). For such complex traits marker assisted selection (MAS) based on a few loci is not expected to significantly increase genetic gain. A newer alternative to MAS is GS (Meuwissen et al., 2001). GS uses all molecular markers distributed across the genome in a regression model to calculate genomic estimated breeding values (GEBVs) of individuals without any prior knowledge of where causal genes are located (Meuwissen et al., 2001). A popular method of implementing genomic prediction (GP) is the genomic best linear unbiased predictor (GBLUP) that utilizes genomic relationships between individuals for prediction. An assumption for the GBLUP model is equal variance for all markers, which is very seldom the case even for complex traits (VanRaden, 2008). Alternatively, identified QTLs can be used to reduce the number of markers expected to influence a trait by combining the GWAS top significant SNPs with a genomic prediction method that allows markers to have different effect sizes. The self-trained and fast nature of machine learning algorithms make them excellent alternatives to traditional genomic prediction methods. Among the most popular machine learning algorithms is random forest (RF), which uses ensemble learning methods of individual decision trees to make an accurate prediction model (Breiman, 2001).

Previous studies have shown that 80-200 GWAS selected markers can predict yield in wheat with an accuracy similar to GP models based on all available marker data (Cericola et al., 2017). However, only few studies have used GWAS to identify QTLs in white clover and none of these examined biomass yield, the genetics of which remains poorly understood (Inostroza et al., 2018; Kaur et al., 2017).

In this study, we examine binary interactions of rhizobium strains and clover genotypes by continuously monitoring plant growth and quantifying dry matter yield. We use these results to assess the relative contributions of clover and rhizobium genetic variation to yield, identify yield-related QTLs and predict yield within and across generations.

## Results

### 145 diverse white clover genotypes derived from commercial cultivars

We carried out RNA-sequencing of roots for a panel of 145 white clover genotypes derived from 20 commercial cultivars and identified 383,280 high quality SNPs that were used for all downstream analyses. The genetic structure of the white clover population was assessed using a genomic relationship matrix (**Figure 1B**). In general, the genotypes clustered by cultivar and showed low levels of relatedness. Genotypes from different cultivars had relationship coefficients close to 0. In line with these results, principal component analysis indicated that the clover genotypes clustered mostly by variety (**Figure 1C**). We found a rapid decline in LD after 1 kb (**Figure 1D**) and high levels of heterozygosity with an average of 0.28 (**Figure 1E**), consistent with obligate outcrossing.

### Initial plant size is strongly correlated with yield

To evaluate the relative contributions of clover and rhizobium genetic variation to white clover yield, we combined the 145 clover genotypes with 169 previously described *Rhizobium leguminosarum* sv. *trifolii* strains representing genospecies A, B, and C (Cavassim et al., 2020). We tested 704 clover-rhizobium combinations in a greenhouse setting, continuously monitoring plant growth using a high-throughput imaging system (Tausen et al., 2020) (**Figure 2A-C**). Using stolon cuttings to generate clones of single plants, we included an average of 15.9 replicates of each white clover genotype in combination with 4-6 different rhizobium strains or a mix of multiple strains, which yielded 13.6 replicates of each rhizobium strain and a total dataset of 2,304 observations. The experiment was structured into two different rounds of a randomised trial design. Each round consisted of two sets where each set refers to a full set of 704 unique clover and rhizobium combinations, each grown in 2-3 replicates per round. The 19 non-inoculated controls showed pale yellow leaves, indicative of nitrogen starvation, and very poor growth, showing that the experimental setup efficiently prevented spread of rhizobia between pots.

**Figure 2.**
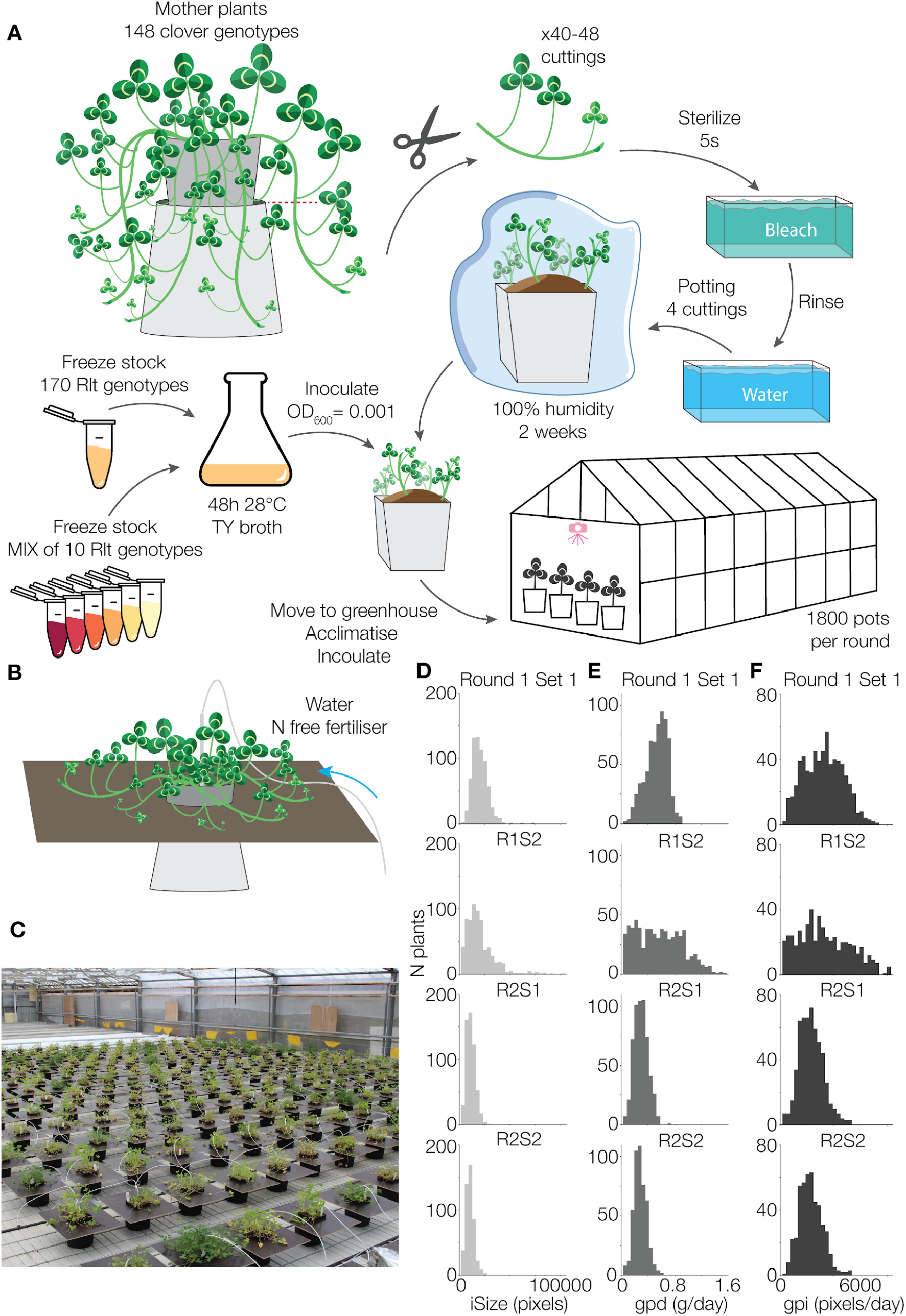
Experimental setup. **A:** Clonal propagation of 148 clover mother plants and the potting method. Each pot was inoculated with either one of 170 individual, characterised *Rlt* strains or a mix of 10 genetically diverse strains. **B:** Individual pot setup and drip watering system. **C:** Picture from the greenhouse showing the setup. **D-F**: Histograms of raw data after filtering for **D:** initial size (iSize, *n* = 2392), **E:** growth per day (gpd, *n* = 2392), and **F:** growth rate during nitrogen fixation (gpi, *n* = 2203).

We recorded dry matter yield as an end-point measurement and calculated the average growth per day (gpd) as the dry matter increase per day from inoculation to harvest (**Figure 2E**). Based on the image data, we quantified the initial plant size prior to onset of nitrogen fixation (iSize) and growth rate during nitrogen fixation (gpi) (**Figure 2D+2F, Supplementary file 1**). iSize was calculated as the average plant size during the first 10 days post inoculation, whereas gpi represented the average growth rate from day 11 to 25 post inoculation (**Figure 3A**). Observations of iSize, gpd and gpi were approximately normally distributed for most combinations of rounds and sets. However, this was not the case for gpd observations from round 1 set 2, where plants had been inoculated on different days and greenhouse temperatures had been unusually high (**Figure 2D-F**).

**Figure 3.**
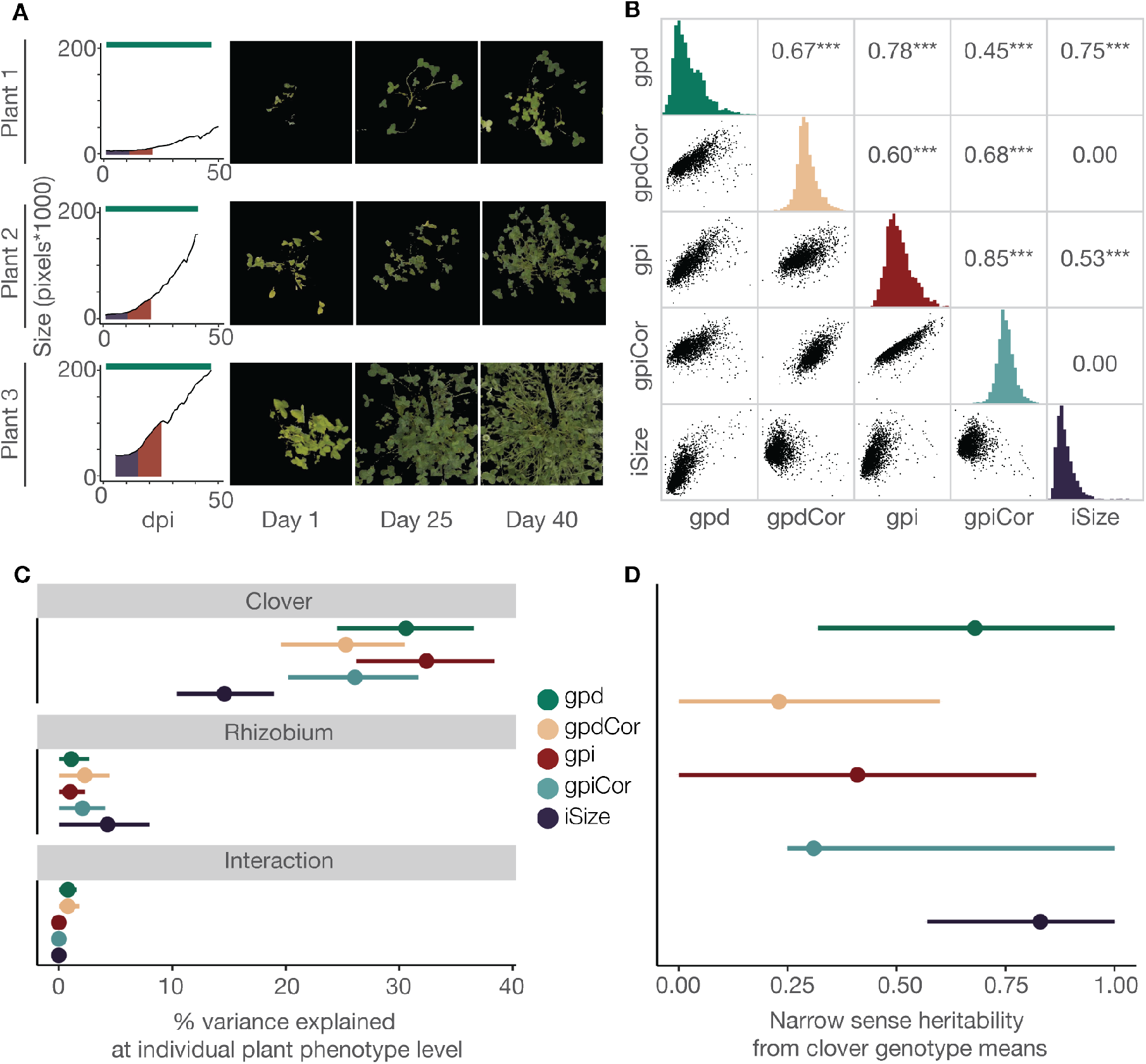
Phenotypic data and trait heritabilities. **A:** Growth curves and masks of individual plants from the clover genotype Banna_0204 at the end of the 10 day initial size interval, after 25 days, and after 40 days. Growth curves indicate the days past inoculation (dpi) used to extract the different phenotypes; day 1-10: iSize (purple), day 11-20: gpi (red), and day 0-harvest: gpd (teal). **B:** Pairwise correlation of traits. The diagonal shows histogram of each trait. Left of the diagonal: scatter plots of pairwise comparisons between gpd, gpdCor, gpi, gpiCor, and iSize. Right of the diagonal: Pearson product moment correlation coefficients. *n* = 2304. **C-D:** Estimation of variance components and heritabilities of different yield-related traits. Estimates (dots) and their 95% highest posterior density intervals (lines) are coloured by trait. **C:** The estimated percentage of phenotypic variance explained by clover, rhizobium or the clover x rhizobium interaction using all observations, *n* = 2304. **D:** The estimated narrow sense heritability (*h^2^*) of the traits when averaging across clover genotypes, *n* = 145. Note that (**C**) is based on individual plant phenotypes, whereas (**D**) is based on clover genotype means. This is why the narrow sense heritability is larger than the broad sense heritability for some traits.

Individual plants of the same genotype showed large variation in both iSize and their overall growth (**Figure 3A**). Based on all 2304 observations, we found that iSize was strongly correlated with gpd (**Figure 3B, Supplementary file 2**). Likewise, iSize and gpi were also significantly correlated, although they represent non-overlapping growth stages (**Figure 3B, Supplementary file 2**). This suggested that variation in plant size prior to symbiosis establishment, iSize, had a large impact on the entire growth period. To correct for this variation, and get independent representations of growth stages prior to and during nitrogen fixation, we subtracted the effect of iSize from gpd and gpi, to obtain the traits gpdCor and gpiCor, respectively.

### Rhizobium variation does not significantly affect yield

Based on the full set of data, which included 3-4 four replicates of each clover-rhizobium combination, we estimated the variance explained by clover, rhizobium and clover x rhizobium interaction for gpd, gpdCor, gpi, gpiCor and iSize. For all traits, we found that clover explained more of the yield variance than rhizobium. The rhizobium contribution was not significant for any of the traits, as the highest posterior density interval (HDPI) included zero (**Figure 3C, Supplementary table 1**). Although significant for gpd and gpdCor, the clover x rhizobium interaction explained a very small part of the variation (**Figure 3C, Supplementary table 1**). The proportion of variance explained by clover was smallest for iSize (**Figure 3C, Supplementary table 1**), which is consistent with considerable stochastic variation in cutting size (**Figure 3A**). On the other hand, correcting for iSize led to a decreased proportion of variance explained by clover for both gpd and gpi, suggesting that iSize includes a genetic component relevant for yield. Based on this broad-sense heritability analysis, we chose to focus exclusively on the clover contribution for the remaining analyses and averaged the data by clover genotype to obtain 145 observations of each trait (**Supplementary file 3**).

### Yield and initial plant size show high narrow sense heritabilities

Using the averaged data, we then estimated narrow sense heritabilities for all traits using a GRM based on the 383,280 RNA-seq SNPs (**Supplementary file 4)**. In contrast to its low broad sense heritability at the single plant level, iSize showed a high narrow sense heritability of 0.83 for the data averaged by clover genotype (**Figure 3D**). This indicates that stochastic variation in cutting size is efficiently controlled for by averaging across a large number of replicates, clearly revealing a large genetic component captured by the genotyped SNPs. Gpd showed the second highest SNP heritability, whereas the remaining yield traits, gpdCor, gpi and gpiCor, which describe the second part of the growth phase, showed lower narrow-sense heritabilities (**Figure 3D**).

The high narrow sense heritabilities for iSize and gpd were encouraging for genomic prediction, and we applied GBLUP prediction models to all traits using 6-fold cross-validation repeated 100 times. We observed moderate to high prediction accuracies for gpd (0.39) and iSize (0.53) **(Figure 4A)**. Further, the model was able to predict gpi with a low accuracy of 0.16 **(Figure 4A)**. However, we were unable to predict the traits where iSize effects were eliminated, gpdCor and gpiCor, from genetic data **(Figure 4A)**.

**Figure 4.**
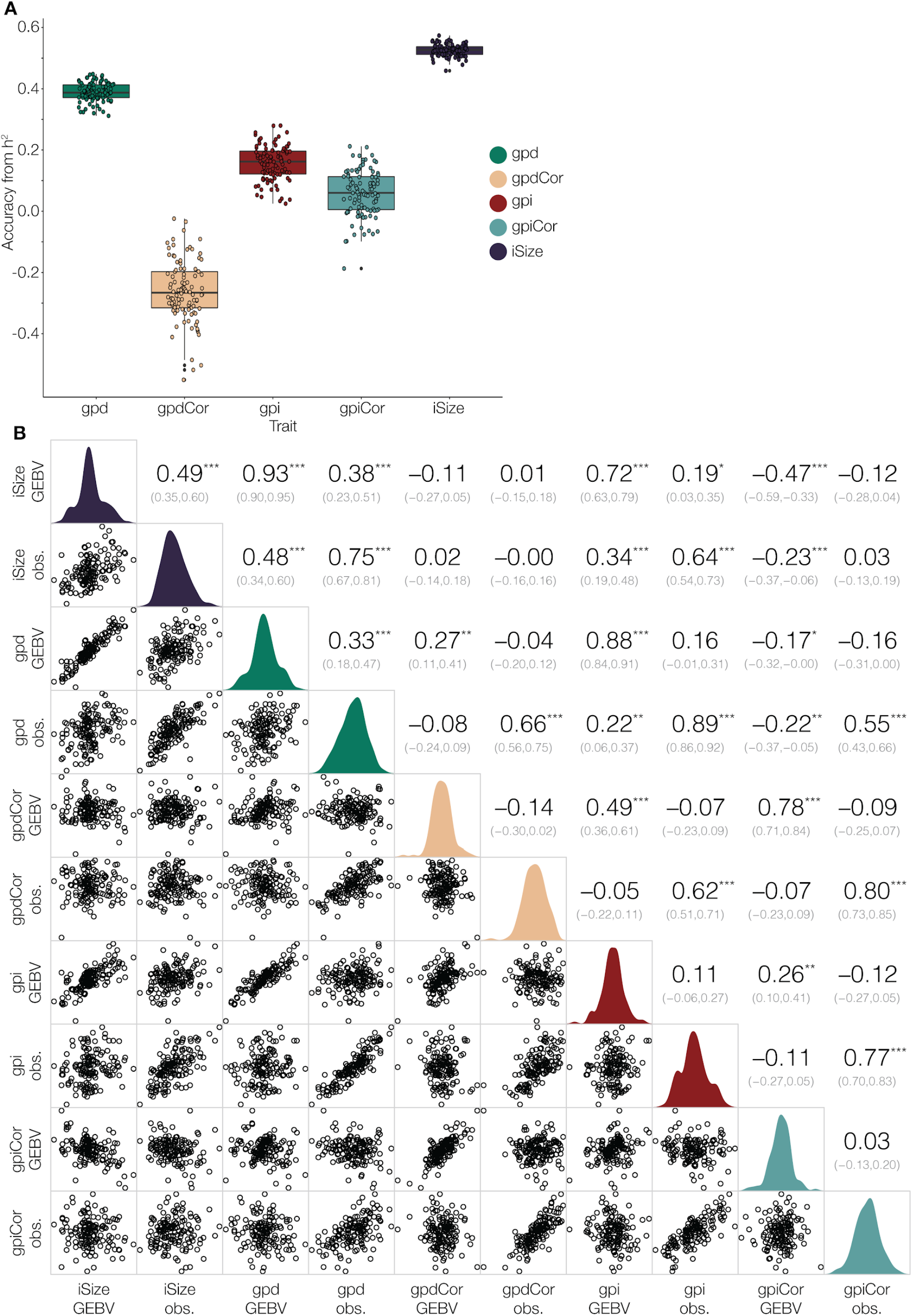
Genomic prediction of gpd, gpdCor, gpi, gpiCor, and iSize. **A:** Boxplot of prediction accuracies based on 100 rounds of GBLUP. **B:** Correlations of traits and their GBLUP-produced GEBVs. Confidence intervals of correlation coefficients are given in parentheses and asterisks indicate significance of correlations *, *p* < 0.05; **, *p* < 0.01; ***, *p* < 0.001. *n* =145.

To test more thoroughly if later growth stages could contribute genetic information relevant to yield, we predicted gpd by combining iSize GEBVs with gpdCor, gpi or gpiCor GEBVs. We found that gpdCor GEBVs could not explain a significant part of the gpd phenotypic variance, whereas gpi and gpiCor GEBVs could explain a significant part of gpd variance only when iSize GEBVs were not included in the model **(Supplementary table 2)**. Based on these results, we focused on the iSize and gpd traits for the remaining analyses.

### Yield can be predicted based on the genetics of initial plant size

To further investigate the relationship between the traits, we calculated Pearson correlation coefficients for all possible combinations of observed phenotypes and estimated GEBVs **(Figure 4B)**. Surprisingly, we observed that gpd showed a higher correlation with iSize

GEBVs (0.38) than with its own gpd GEBVs (0.33). Furthermore, we found the correlation of GEBVs of gpd and iSize to be 0.93, indicating that the two traits show a very high degree of genetic correlation that exceeds the phenotypic correlation of 0.75 (**Figure 4B**). These observations indicate that it is possible to predict gpd from iSize, i.e. to predict dry matter accumulation for the entire growth period based on GEBVs obtained exclusively from data describing the initial growth phase prior to onset of nitrogen fixation.

Since iSize could be a comparatively simple trait, if for instance related mainly to leaf size, we carried out GWAS to identify specific markers associated with iSize and/or gpd. In line with the high level of genetic correlation, we observed overlapping genetic signals associated with gpd and iSize on chromosomes 3 and 7 (**Figure 5A-D, Supplementary figure 1, Supplementary file 5**). In addition, we found a strong signal approaching the Bonferroni-threshold on chromosome 13 and a peak on chromosome 1, which was exclusively associated with iSize. In general, the gpd associations were weaker than those of iSize, but we found a gpd signal on chromosome 8 that was not identified for iSize (**Figure 5A-D, Supplementary figure 1)**.

**Figure 5.**
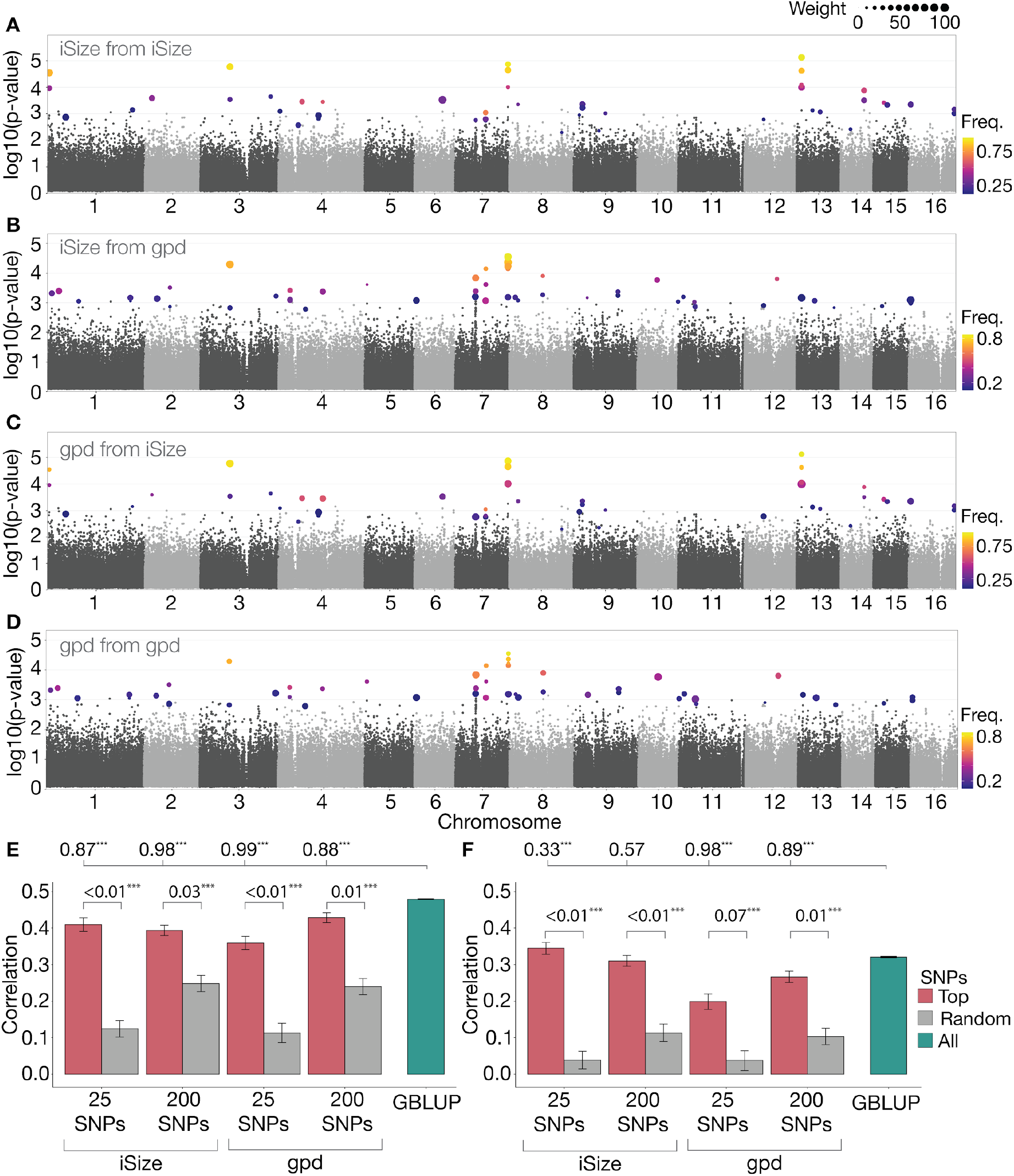
GWAS and genomic prediction using top GWAS markers. **A-D:** Manhattan plots coloured by how often a given SNP occurs in the 25 most significant GWAS SNPs and scaled by the predictive importance given by the RF algorithm. Frequency of occurrence (Freq.) is calculated as a fraction of the 600 times a GWAS was run. Only SNPs with a top 25 frequency > 0.10 were coloured. **A:** Prediction of iSize based on iSize GWAS. **B:** Prediction of iSize based on gpd GWAS. **C:** Prediction of gpd based on iSize GWAS. **D:** Prediction of gpd from gpd GWAS. **E-F:** Correlation between predicted GEBVs and observed phenotypes when using different methods for prediction of iSize **(E)** or gpd **(F)**. The red bars display the correlation obtained when using the top 25 or 200 most significant SNPs obtained by performing GWAS using either iSize or gpd. The trait used for GWAS is specified below the *x* axis. The blue bars display the correlation obtained using GBLUP on all markers. Grey bars display the correlation obtained when using 25 or 200 random SNPs as input in the RF model. Error bars display standard errors obtained when repeating the experiment 100 times with 100 different divisions into test and training populations. The numbers at the very top of the plot indicate the fraction of times out of 100 that the given method was outperformed by the GBLUP method. The asterisks following the fractions indicate whether the model performed differently from GBLUP according to a paired sample sign test. ***, *p* < 0.001. The numbers comparing the red and grey bars indicate the fraction of times out of 100 that the method built on random SNPs outperformed the method built on top GWAS SNPs. The asterisks following the fractions indicate whether the top SNPs performed differently from random SNPs according to a paired sample sign test. ***, *p* < 0.001.

Motivated by what appeared to be clear GWAS signals, although they did not reach genome-wide significance, we set up a two-step prediction approach. First, we conducted a GWAS using a training population to select the top 25 or 200 most significant markers. Second, we used a random forest (RF) approach to predict gpd or iSize in a testing population based on the GWAS markers. GWAS was carried out for 600 different training populations, resulting in 949 unique SNPs in top 25 for iSize and 1196 for gpd. When considering only SNPs that occurred in at least 10% of the GWAS runs, the numbers were reduced to 43 and 47 for iSize and gpd, respectively. We predicted iSize and gpd based on both iSize and gpd GWAS markers. All GWAS-based predictions were compared to predictions using 25 or 200 random SNPs. The predictions based on the top GWAS SNPs were significantly more accurate than those based on random sets of markers in all cases (**Figure 5E+F**).

For iSize, the GWAS+RF method achieved performances close to GBLUP using the full GRM, but the GBLUP model was significantly better in all cases **(Figure 5A)**. For gpd, the predictive performance was relatively poor when using the top 25 or top 200 gpd GWAS SNPs for prediction. Especially, using the top 25 gpd SNPs resulted in a large drop in predictive power, with an average correlation of 0.20 compared to 0.33 using GBLUP. The iSize-associated markers produced more accurate predictions for gpd, indicating that the iSize trait more accurately captures the relevant genetics. Predicting gpd from the top 200 iSize GWAS SNPs did not differ significantly from the GBLUP results. However, using the top 25 iSize markers resulted in significantly better accuracy than GBLUP (**Figure 5F**).

To evaluate the stability and importance of the genetic regions associated with gpd and iSize, we coloured markers by the fraction of times they occurred in the top 25 most significant SNPs in our cross-validation scheme and scaled them by their average importance given by the RF model (**Figure 5A-D)**. The top SNPs contributing the overlapping iSize/gpd peaks on chromosomes 3 and 7 occurred frequently in top 25, thus providing us with stable and trustworthy signals for both traits (**Figure 5A-D)**. The SNPs in the remaining peaks identified for iSize also showed high occurrences **(Figure 5A+C)**. The SNPs in the peak on chromosome 8 identified exclusively in the gpd GWAS showed relatively low occurrence, indicating that they were less likely to be truly associated with gpd (**Figure 5B+D**). Looking further into the prediction of gpd from the top iSize markers and *vice versa* we found that predicting a trait based on GWAS results from the other correlated trait caused the genetic regions shared by the two traits to be assigned more importance by the RF model relative to the peaks specific for one trait (**Figure 5A-D).** Overall, the GWAS and RF prediction results indicated that relatively few genomic regions contribute large effects to the traits of interest. The genes located most closely to the markers in the GWAS peaks on chromosomes 1, 3, 6, 7, 8 and 13 are listed in **Supplementary table 3**. These include a putative ortholog of *Arabidopsis thaliana GIGANTUS1*, which regulates growth and biomass accumulation (Gachomo et al., 2014).

### F1 poly-cross yield can be predicted with high accuracy

The greenhouse experiment was based on stolon cuttings in order to obtain genetically identical individuals in an outcrossing species. However, white clover is normally grown from seed. To examine the relevance of our data for seed-grown plants and to evaluate the ability to predict performance across generations, we set up nine poly-crosses with 4-6 parents chosen among the 145 genotypes tested in the greenhouse (**Figure 6A, Supplementary table 5**). For both iSize and gpd, the parents represented a large diversity in their GEBVs as predicted by GBLUP (**Supplementary figure 2, Supplementary table 5**). The F1 populations resulting from the polycrosses showed large differences in average yield, ranging from 2.9 g to 4.6 g of dry matter per F1 individual (**Figure 6B, Supplementary table 5, Supplementary file 6**).

**Figure 6.**
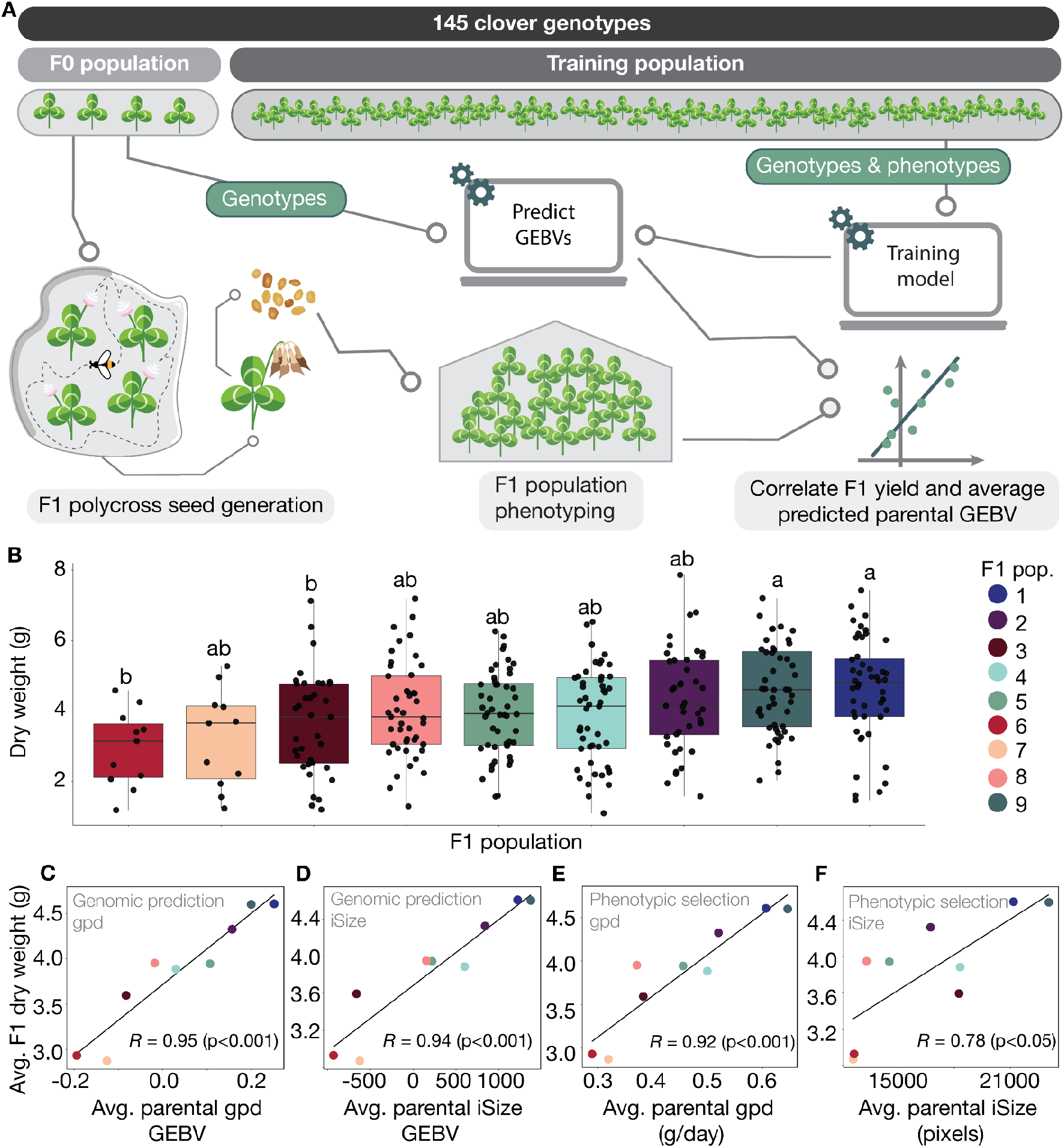
F1 crossing and genomic prediction strategy. **A:** Experimental and prediction setup. **B:** Boxplot for F1 phenotypic data for the 9 polycrosses. F1 population means with different letters differ at *p* < 0.05 according to Tukey honestly significant difference (HSD). **C-F:** F1 cross generation prediction. **C-D:** Correlation between average F1 population yield and average GEBVs of gpd (**C**) or iSize (**D**). **E-F:** Correlation between average F1 population yield and average parental phenotypes of gpd (**E**) or iSize (**F**).

To test if we could predict the average yield of the nine F1 populations based on the data from the parental (F0) generation, we calculated GEBVs for the F0 parents, using the remaining 139 to 142 F0 clover genotypes as the training population (**Figure 6A**). This simulates a scenario, where the parents for a synthetic cultivar are genotyped but not phenotyped, and the average genotype of an offspring population is assumed to be represented by the average parental GEBV. We found a correlation between the average parental GEBV and the average dry weight of F1 populations of 0.95 for gpd and 0.94 for iSize (**Figure 6C-D)**. We compared these values to the correlation between F1 average dry weight and the average F0 phenotype. This represents a phenotypic selection scheme, where parents are selected directly based on their F0 gpd or iSize phenotype. Phenotypic selection yielded correlations of 0.92 and 0.78 for gpd and iSize, respectively (**Figure 6E-F**). We conclude that genomic prediction performed better than or equal to direct phenotypic selection. In particular, genomic selection outperformed phenotypic selection for iSize, indicating that it is not initial plant size *per se*, but rather its heritable genetic components that are important for yield.

## Discussion

Previous studies on white clover-rhizobium effects on yield relied on a handful of clover and rhizobium genotypes (Mytton, 1975; Mette M. Svenning et al., 1991; Young & Mytton, 1983), likely because the pairwise testing of many clover-rhizobium combinations requires a large experimental setup. Here, we examined more than 700 different combinations, but found no significant contribution to yield from rhizobium and only minor effects of clover-rhizobium interactions. Since we used diverse rhizobium strains belonging to three distinct genospecies of *R. leguminosarum* sv. *trifolii*, which were collected from locations in Denmark, England and France (Cavassim et al., 2020; Moeskjær et al., 2020), we had initially expected variation in nitrogen fixation efficiency between the strains. All strains were collected from pink and healthy looking root nodules, suggesting an effective symbiotic interaction. In our setup, this generalised to efficient interactions with all tested clover genotypes, although they too represent considerable genetic variation across twenty different commercial cultivars. Our results suggest that white clover is efficient in sanctioning rhizobia that do not provide high levels of fixed nitrogen and that a large degree of cross-compatibility exists for European white clover and *R. leguminosarum* sv. *trifolii* genotypes. This does not rule out that different clover genotypes would preferentially select specific rhizobium strains or that strong clover-rhizobium interaction effects exist outside of our population samples, as reported by others (Irisarri et al., 2019). Indeed, if we had also isolated rhizobium strains from small and inefficient-looking nodules, we may well have identified such effects. Seen from an inoculation perspective, our results indicate that efficient rhizobia are present and selected by white clover at all sampled locations, and our isolated strains represent a genetically well-characterised source of potential new white clover inoculants for locations with limited rhizobium populations.

We used stolon cuttings to achieve genetic replication in an outbreeding species. Because of the insignificant contributions from rhizobium, we effectively had more than ten replicates of each clover genotype. This helped reduce the effects of variation in cutting size and allowed us more accurate phenotype estimates for each clover genotype. For F0 predictions, reducing the number of replicates quickly leads to deterioration of prediction accuracy, whereas F1 polycross predictions retain high accuracy even with as little as two replicates (**Supplementary figure 3**). Because the polycrosses have 4-6 parents, however, two replicates in the F1 predictions correspond to averaging across at least eight data points. The prediction of polycross yield is therefore intrinsically more robust to limited replication of individual genotypes. Still, accurate predictions of the breeding values of potential parents is required for ensuring maximum genetic gain, and here increasing the number of replicates, even to more than 10, appears to result in increased accuracy (**Supplementary figure 3**). Clonal propagation is used in many major food crops including nearly all types of fruit and important roots and tubers, making it highly relevant to consider the effect of the number of replicates on prediction accuracy (Bradshaw, 2016; Grüneberg et al., 2009). It is worth noting that the stolon-based greenhouse experiments were carried out in two rounds, which differed both with respect to growth medium (sterilized peat or vermiculite) and time of year (spring or summer). Because of this large variation in environmental conditions, we consider it likely that the genetic associations discovered using the complete data set are generally important for white clover yield potential. The fact that yield from seed-grown F1 plants cultivated the following year in a different greenhouse could be predicted with high accuracy based on the stolon cutting data supports this hypothesis.

We find it striking that we could predict yield most accurately based on the genetics underlying initial plant size from day 1-10 post inoculation, and that data from the remaining growth period, even the dry matter yield data itself, did not contribute additional relevant genetic information. Others have reported genetic correlations between growth at different stages and biomass yield in rye, but the relative predictive power of the different observation was not investigated in detail (Miedaner et al. 2018). The image data was critical for understanding the characteristics and limiting factors for yield, because it enabled us to examine distinct growth stages separately. Under our experimental conditions, the clover yield potential was already manifest in the size of the stolon cuttings from fully nitrogen-fertilized mother plants, indicating that variation in nitrogen fixation efficiency in later growth stages did not impact yield. This is consistent with the lack of substantial contributions from rhizobium and clover-rhizobium interactions. In contrast, iSize captures critical yield components related to morphology, probably most prominently leaf size. In field trials, leaf size was previously identified as a trait with very high narrow sense heritability, while dry matter yield showed moderate heritability and the two traits displayed a positive genetic correlation (P. Annicchiarico et al., 1999; Jahufer et al., 1994).

Previous studies used pedigrees to estimate narrow sense heritability, whereas we base our analysis on material genotyped using RNA-seq, which yields a large number of markers located within genes. This allowed us to identify specific candidate genes associated with yield and initial plant size. White clover has complex allotetraploid genetics, and a large number of densely distributed markers is required to detect signals linked to causal loci because of the very low LD in outbreeding population. QTLs associated with white clover cold tolerance were identified using a tetraploid model, whereas no signals could be detected using a diploid model (Inostroza et al., 2018). However, Inostroza et al. did tetraploid genotype calling which gave a different starting point for diploid GWAS than in our models. When calling the genotypes as diploid, we found that using a standard diploid GWAS worked well. We would expect that to be the case since the two white clover sub genomes exist in parallel without recombining (Griffiths et al., 2019), which allows us to assume the presence of only two alleles per locus. It is intriguing that one of the strong GWAS signals was located near a homolog of Arabidopsis *GIGANTUS1*, which regulates biomass accumulation, potentially through effects on ribosome biogenesis. In fact, the top markers on chromosomes 3 and 7 appear very tightly linked (**Supplementary figure 4**). One of the peaks is likely misplaced and the two signals probably represent a single peak near the *GIGANTUS1* gene. It has not been studied in biomass crops, and further studies will be required to determine if there is a causal effect of *GIGANTUS1* variation with respect to white clover biomass yield. The results of our cross validation scheme for prediction using GWAS SNPs in an RF model provides compelling evidence for the predictive value of the associated markers. Since we were able to predict yield more accurately using the top 25 GWAS markers than using a GRM based on the full set of SNPs, marker-assisted selection could potentially yield quick progress simply by applying it to subselection within existing cultivars, none of which are fixed for the alleles with positive effects on yield or initial plant size (**Supplementary figure 4**). Our finding that we can predict yield well using a relatively small set of markers is quite surprising. Compared to an inbreeding crop like wheat, white clover has high levels of heterozygosity and rapid LD decay, which should make it difficult to select small marker-sets for prediction. In wheat, 200-300 markers were needed to achieve a prediction accuracy similar to that using GBLUP (Cericola et al., 2017; Inostroza et al., 2018). This suggests that we have successfully identified major yield QTL in white clover. Likewise, it was encouraging to see the very high prediction accuracy for F1, indicating that genomic prediction can be a very robust tool for prediction of polycross performance.

The main limitations of the study is that our experiments were carried out in the greenhouse in order to be able to control the rhizobium populations and in the absence of a companion grass. The greenhouse is a warm and well-watered environment that to some degree shelters the plants from the environment, and additional factors will certainly affect field grown material. However, it is promising to see specific markers strongly linked to yield and plant size, and future trials will tell if these remain relevant in the field. Considering the results presented here, it is tempting to reiterate the suggestion to base white clover breeding on well-replicated cloned material, deferring progeny testing to late stages in the selection programme (P. Annicchiarico et al., 1999; Gibson et al., 1963). Despite a larger initial investment, determining the genetic merit of individual plants using cloned plants might be what is needed to significantly accelerate genetic gains in forage legumes.

## Materials and Methods

### Plant material and clonal propagation

The plant material used in this study consists of a panel of 148 white clover genotypes from 20 commercial varieties with diverse agronomic qualities. To cover maximum genetic diversity, the genotypes were chosen to be as morphologically distinct as possible within each variety. To ensure genetically identical replicates from each genotype, individual plants were clonally propagated from mother plants. Four stolon cuttings were taken from the mother plants with sterilised scissors including a minimum of three internodes and viable leaves. The stolons were sterilised in 1:4 bleach (Klorin, Colgate-Palmolive Company, USA) for five seconds, washed with tap water, and subsequently stored in tap water with Conserve (Dow Agroscience, Denmark) until potting (1 to 10 minutes) to prevent transfer of thrips from our breeding greenhouse to the experimental greenhouse. An overview of the experimental setup can be seen in **Figure 2A** and **2C.**

### Greenhouse setup

The plants were grown in individual 5L pots under greenhouse conditions in Egå, Denmark (56.226°N, 10.259°W). Water and nutrients were supplied through individual feeding tubes to minimize contamination between pots (**Figure 2B**). The experiment was structured into two rounds of a randomised trial design with 10-24 replicates per clover genotype. Each round consisted of two sets where each set refers to a full setup of 883 unique clover and rhizobium combinations, each grown in 2-3 replicates per round. In round 1, cuttings were potted in gamma irradiated peat (Pindstrup Mosebrug A/S, Denmark) (**Supplementary table 4**) and stored under white plastic at 100% humidity for two weeks. The growth periods of Round 1 Set 1 and Set 2 were from 11/05/2018 to 02/07/2018 (52 days) and from 05/06/2018 to 26/07/2018 (42 to 49 days), respectively. In round 2, plants were potted in vermiculite (Pull Rhenen B.V., Netherlands) and stored under white plastic at 100% humidity for two weeks. The plastic storage sacks containing vermiculite were sterilised with Klorin prior to opening. The growth period of both sets in Round 2 was from 15/08/2018 to 24/10/2018 (68 to 70 days). All plants were acclimatised for a week and transferred to the main greenhouse for the trial. Immediately after removing the plastic, plants were inoculated with one of 169 genetically characterised *Rhizobium leguminosarum* bv. *trifolii* strains (Cavassim et al., 2020), or a mix of 10 genetically distinct strains (OD_600_ = 0.001). One genotype (Aearl_07) was highly replicated and inoculated with all strains, separately. Only pots where four cuttings survived until the end of the trial were included in the analyses. Further, observations from uninoculated plants or plants inoculated with ‘SM73’ were removed due to contamination. To avoid a large contribution from the many Aearl_07 observations to the subsequent analysis, its observations were scaled down to represent only observations from plants inoculated with six random symbionts. The detailed experimental setup and overview of the greenhouse is available in **Figure 2A-C**.

Plants were harvested at the end of the growth period. Harvested plant material was dried at 35°C until a constant weight was achieved.

### Image-based filtering

Plants were monitored using a Raspberry Pi based imaging setup (Tausen et al., 2020). The imaging setup gave rise to daily area measurements of individual plants that were used to estimate the initial size of single plants expressed as the average size of the plant during the first 10 days of growth using the Greenotyper software (Tausen et al., 2020). In addition, the daily area measurements of single plants were used to clean up the data by removing presumably unsuccessfully inoculated plants and error-prone measurements. To identify problematic data, area measurements of single plants were regressed on days past inoculation (dpi). Plants with regression coefficient < 100 area/day in the interval 10-20 dpi were considered to be unsuccessfully inoculated and consequently removed. Furthermore, plants that showed an overall negative regression coefficient from 10 dpi to the remaining growth period were removed. In total this image-based filtering removed data points of 163 plants. In addition, we removed 3 clover accessions we did not have genotype data for. This gave a total of 2304 observations from 704 combinations of 145 clover and 169 rhizobium genotypes including 6-20 replicates pr. clover genotype.

### Genomic data

RNA from a panel of 148 white clover accessions was sequenced using Illumina 150 bp Paired End reads (Novogene, Hong Kong). The RNA used for genotyping was extracted from roots of plants grown in sterile vermiculite for 8 weeks. Roots were washed with sterile water, harvested, and immediately frozen in liquid nitrogen. 2mm of the root tip was removed prior to freezing. RNA was isolated using the NucleoSpin RNA Plant (Macherey Nagel, Germany).

RNA-seq reads were mapped to the reference S9 genome using the STAR software (v2.5.2) with stringent mapping to avoid ambiguous mapping between the two subgenomes (Dobin & Gingeras, 2016). Variant calling was performed for each sample separately using the HaplotypeCaller program in GATK (v3.8) outputting all confidently callable sites (McKenna et al., 2010). The outputs were then merged in batches of 20 using CombineGVCFs and finally combined into a single GVCF file which was genotyped using GenotypeGVCFs. Using SelectVariants in GATK the following filters were applied to the .vcf file: mapping quality > 30, depth > 160 and quality > 20. Annotation was done using bcftools (Danecek & McCarthy, 2017). The workflow for mapping and variant calling can be found at: https://github.com/MarniTausen/WhiteCloverRNAseq.

A number of filtering steps were applied to the raw variant data based on the S9 reference calling: all variants that are not classed as single nucleotide polymorphisms (deletions, insertions, etc.) were excluded, read depth > 300, excess heterozygosity (Phred-scaled *p* value for exact test of excess heterozygosity) < 150, and AN (Total number of alleles in called genotypes) > 130. SNPs with more than 10% missingness were excluded, and the remaining missing SNPs were imputed using BEAGLE version 5 with default settings (Browning et al., 2018). Further, markers with a minor allele frequency < 5% were removed and an LD-filter was applied to remove redundant information by filtering out SNPs that showed complete LD with SNPs already in the data set. In addition, SNPs that showed < 0.5 correlation with genotypes of all other SNPs located within the same gene/intergenic region were removed, as these were considered unreliable. The final set of markers consisted of 383,280 SNPs.

### Population structure analysis

The genomic relationship matrix (GRM) for the clover genotypes was calculated as proposed by VanRaden method 1 (VanRaden, 2008):

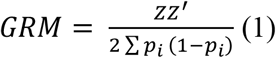

Where *Z* is the centered genotype matrix with dimensions *n* x *m*, where *n* is the number of individuals and *m* is the number of markers. *p_i_* denotes the allele frequency of the second allele at locus *i*. After closer investigation of the GRM, 3 individuals were removed, as they showed very close relationship to an already present sample, and no relationship to the remaining accessions of the variety that they were labeled as belonging to, indicating a labelling error that could not be untangled. All analyses were therefore based on 145 unique genotypes.

Based on the GRM, a principal component analysis (PCA) was performed using the prcomp R-function and the ggfortify package for visualisation (Tang et al., 2016; R Core Team, 2020).

### Multiparental crosses

10 F1 populations were generated from plants in the 145 clover genotype greenhouse setup (**Supplementary table 5**). Crosspollination was done using bumble bees in net houses. Between 20 to 48 F1 seeds were germinated based on available seedstock for each population and grown under greenhouse conditions. Seeds were scarified using sandpaper, germinated for 7 days in petri dishes, and transferred to 0.5 L pots with sterile vermiculite. All plants were inoculated with the same *Rhizobium* strain (SM42). A table watering system with the same fertiliser solution as described in the section “Greenhouse setup and phenotyping” was used throughout the growth period. After 98 days of growth under artificial light, the plants were harvested, dried, and weighed using the approach described above. One of the 10 F1 populations was excluded from the downstream analysis due poor germination and/or growth resulting in < 10 offspring plants. The remaining populations had between 11 and 48 data points. Further, observations within each population with a dry weight below 1g or fresh weight below 10g were removed, since these plants had established poorly and appeared wilted.

### Traits

Initial size (iSize) was measured by pixel counts of a plant from a 512 x 512 pixel mask in the first 10 days of growth after inoculation, i.e. before the symbiotic relationship between the plants and rhizobia strains is established. Another measurement for yield was gpd, which was reported as the dry weight of harvested plants divided by days of growth from inoculation to harvest. For this reason, gpd was overlapping with the iSize measure.

To get a yield measure that was less correlated with the iSize of plants, we calculated three additional yield traits: gpdCor, gpi and gpiCor.

gpdCor reports gpd corrected for the full effect of iSize.

The following equations were applied:

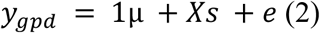

Where *y_gpd_* reports the observed gpd values, *μ* is the intercept, *s* is the fixed effect of initial size and *e* is a vector of residuals. *X* is a design matrix of *n* x 1 dimension, where *n* is the number of observations with observed initial sizes. Estimates from equation 1 was then used to calculate gpdCor:

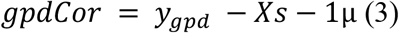

Where variables and matrices are the same as reported in (2).

The fitting of initial size as a fixed effect was done using the “lme4” package in R (Bates et al., 2015).

Growth post inoculation (gpi) reports the growth per day during day 11 to day 25 past the inoculation date. The time interval was set based on a comprehensive test of the trait heritability for growth periods during different time periods and with different lengths. The trait was calculated by using the image data from the greenhouse to fit a regression model to describe the linear relationship between days post inoculation (dpi) and the area of a plant. This can be written as follows:

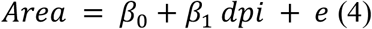

*β*_1_ from equation 3 was then considered our gpi trait. Although plant growth is generally exponential rather than linear, we found the linear regression a good approximation in this growth interval.

gpi was corrected for the full effect of iSize in a similar way as described for gpd in equations 2-3 to produce gpiCor.

### Phenotypic data analyses

The variance estimates of clover, rhizobium and clover x rhizobium interactions were calculated using the following mixed-model on the full data (*n* = 2304).

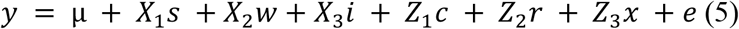

Where *y* is the vector of a trait, *μ* is the overall mean, *s* and *w* are vectors reporting the spatial coordinate of a plant in the greenhouse along the north-south or east-west axis, respectively, *i* is a vector reporting the inoculation date of plants, *c* is a vector of clover effects, *r* is a vector of rhizobium effects, *x* is a vector of clover x rhizobium interaction effects and *e* is the vector of residual effects. *X_n_* and *Z_n_* are design matrices of fixed and random effects, respectively. 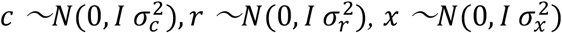 and 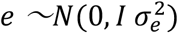 where I is an identity matrix, 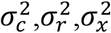 and 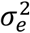 are the variances of clover, rhizobium, clover with rhizobium interaction and the residual effects, respectively.

After this analysis, the average phenotype of a clover genotype was calculated and used for the input in all subsequent analyses (n = 145) including the calculation of the narrow-sense heritability and genomic prediction.

To estimate the narrow sense heritability the following model were fitted:

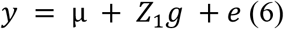

where *y* is a vector of 145 observations corresponding to the average performance of each clover genotype, and *g* denotes a vector of breeding values obtained from the following: 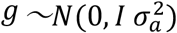 where 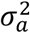 is the additive genetic variance as captured by the GRM. All the remaining terms are as described in equation (5)

The narrow sense heritability was calculated as:

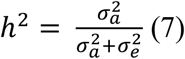

Model parameters were estimated using a Bayesian mixed model relying on a Markov chain Monte Carlo (MCMC) with a length of 20,000 cycles and a burn-in of 5000. The prior distributions were uniform for fixed effects.

This and the estimation of the highest posterior density intervals (HPDIs) was implemented using the BayzR R-package which can be found at: https://github.com/MarniTausen/BayzR.

### Prediction models

Two different approaches were used for genomic prediction of yield-related traits in the population of 145 clover genotypes. These models include a genomic best linear unbiased predictor (GBLUP) model and a two-step method where a genome-wide association study (GWAS) approach is combined with a random forest machine learning algorithm.

In general the GBLUP model can be written as follows:

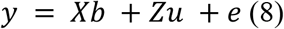

Where *y* is a vector of phenotypes, *b* is a vector of fixed terms which as a minimum includes the overall mean, *u* is a vector of random effects and contains the GEBVs of all genotyped individuals, *e* is the vector of residual effects, and *X* and *Z* are design matrices of fixed and random effects, respectively. 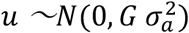 and 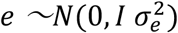 where *G* is the GRM, *I* is an identity matrix, 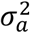 is the additive genetic variance and 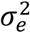 is the residual variance.

Using the GBLUP we modeled the gpd response as follows:

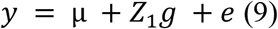

Where *y* is a vector of 145 observations for the yield-related trait, *μ* is the overall mean, *g* is a vector of additive genetic effects from the GRM, and *Z* and *e* are as in (8). The GBLUP model was fitted using the BGLR R-package where the total number of iterations was 20,000 and the burn-in was 5000 (Pérez & de los Campos, 2014).

In the second approach used for yield prediction the first step included a GWAS performed using a python implementation of the EMMAX algorithm followed by *p* value adjustment using EMMA on the top 200 most significant markers (Kang et al., 2008, 2010). The MAC parameter was set to 6. The implementation can be found here: https://app.assembla.com/spaces/atgwas/git/source. Markers were then ordered from lowest to highest *p* values, and a genotype file based on *n* top markers, or *n* random markers were produced. *n* was set to 25 or 200. In the second step this genotype file was used as the input for the RF ML algorithm proposed by Breiman (Breiman, 2001). The RF algorithm was implemented in R using the package “caret” (Max Kuhn, 2020) with the ranger method. The importance of each marker was estimated by using the in-built permutation variable importance approach, which permutes the genotypic values associated with a given marker and then tests the accuracy of the resulting trees and compares it with the accuracy of the tree produced before permutation. The variable importance is then estimated as the difference between the accuracy values and finally scaled to be between 0 and 100 (Wright & Ziegler, 2017; Max Kuhn, 2020).

### Cross-validation

The performance of the prediction models were evaluated using a 6-fold cross validation scheme that was repeated 100 times. In this scheme phenotyped individuals were randomly divided into 6 non-overlapping subsets of similar sizes. Each subset (⅙) then took turns functioning as the testing population by having phenotype values masked and predicted from the phenotypes and genotypes of the remaining (⅚) individuals contributing the training population. In the GWAS+RF method, the testing populations were excluded from the GWAS study, meaning that top SNPs were estimated based on the training population alone. The predictive ability was estimated calculating the Pearson correlation between genomic estimated breeding values (GEBVs) and the observed phenotypes. The significance of a correlation was tested using the agricolae package in R (de Mendiburu, <u>2010</u>).

The prediction accuracy was calculated by dividing the predictive ability with the square root of the narrow sense heritability (equation 9):

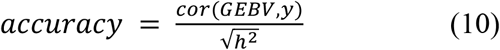

We also set up a validation system to predict yield across generations. This was done by having the 4 to 6 parents of an F1 population constitute the testing population and have the remaining 141 to 139 accessions constitute the training population using their gpd traits and genotypes to train the model. To assess the predictive ability of the cross-generation prediction, Pearson correlations were calculated between the average dry weight and the average parental GEBV of the nine F1 populations, naively assuming that all parents had contributed equally to an F1 population.

### Statistical tests for comparison of prediction methods and F1 means

To test whether prediction methods differed significantly in their predictive ability the following sign test were applied:

First we set up a null hypothesis stating that the predictive ability of method A and method B did not differ in performance. That is on average we would expect the correlation of method A to come out higher than method B in 50% of the cases and lower in the remaining 50% of the cases due to randomness. We viewed the distribution as binomial, calculating the number of successes (*x*) as the observed number of times method A outperformed method B in the 100 repeats (*n*). We then used the inbuilt pbinom function in R to calculate the cumulative probability of x successes or less in *n* observations given a probability of 0.5. This probability was reported as the *p* value. For x > 50 we calculated p as 1 subtracted the cumulative probability of *x* successes in *n* trials given probability 0.5. Consequently, *p* values report the probability of the observed or something more extreme.

The means of F1 population dry weights were compared with a Tukey test in R using the built-in Tukey honestly significant difference (HSD) function.

All scripts used for statistical analyses and visualisation of data is available at: https://github.com/cks2903/White_Clover_GenomicPrediction_2020

### Replicate reduction

Prior to the replicate reduction analyses, the full data set (*n* = 2304) was filtered to include only genotypes with at least 10 replicates which included a total of 142 genotypes. Subsequently, random replicates were removed for each genotype until only 10 replicates were left per genotype. Phenotypes were then averaged for genotypes, and a six-fold cross validation was used to estimate the Pearson correlation between the observed phenotypes of a yield trait and the predicted GEBV. Replicate reduction then followed in a stepwise manner which removed one additional random replicate pr. genotype in each step, calculated the resulting genotype mean, and tested the resulting correlation. The full stepwise reduction was repeated 100 times. The replicate reduction was applied to the cross-generation prediction as well.

## Author Contributions

Conceptualization, S.U.A., S.M; Methodology, C.K.S., S.M., S.U.A., L.J.; Software, C.K.S., L.J., M.T, S.M., R.W.; Validation, C.K.S., M.T., S.M., S.U.A., L.J.; Formal Analysis, C.K.S., S.M., M.T., R.W; Investigation, S.M., C.K.S, M.T., N.R.; Resources, L.J., S.U.A., N.R.; Data Curation, C.K.S., S.M., M.T.; Writing – Original Draft, C.K.S., S.M.; Writing – Review & Editing, S.U.A., C.K.S., S.M.; Visualization, S.M., C.K.S.; Supervision, S.U.A., L.J.; Project Administration, S.U.A.; Funding Acquisition, S.U.A.

## Acknowledgements

We thank Finn Pedersen, Nanna Walther, Mike Ladefoged Damholdt, and Karina A. Kristensen for plant work and greenhouse tending, M. Izabel C. Alves, Trine F. Gadeberg, Caroline Benfeldt, Camous Moslemi, Leandro A. Escobar-Herrera for help in the greenhouse, and Marc Clausen for implementing the technical part of the imaging system. This work was funded by grant no. 4105-00007A from Innovation Fund Denmark (S.U.A.).

## Supplementary tables

**Supplementary table 1.** Estimation of variance components of different yield-related traits. Variance estimates include clover, rhizobium, interaction between clover and rhizobium (Clover x Rhizobium) and residual variance. *n* = 2304.

| Trait | Variance |  |  |  |
| --- | --- | --- | --- | --- |
|  | Clover | Rhizobium | Clover x Rhizobium | Residual |
| **gpd** |  |  |  |  |
| Estimate | 1.2E-02 | 4.2E-04 | 3.2E-04 | 2.6E-02 |
| HPDI | (8.6E-03, 1.5E-02) | (0.0, 1.2E-03) | (2.0E-05, 7.6E-04) | (2.4E-02, 2.8E-02) |
| Variance explained (%) | 30.6 | 1.1 | 0.8 | 67.5 |
| **gpdCor** |  |  |  |  |
| Estimate | 5.4E-03 | 4.8E-04 | 1.8E-04 | 1.5E-02 |
| HPDI | (3.9E-03, 6.9E-03) | (0.0, 1.0E-03) | (2.8E-05, 4.3E-04) | (1.4E-02, 1.6E-02) |
| Variance explained (%) | 25.3 | 2.3 | 0.8 | 71.6 |
| **gpi** |  |  |  |  |
| Estimate | 4.3E+05 | 1.8E+04 | 1.5 | 1.2E+06 |
| HPDI | (4.3E+05, 7.5E+05) | (0.0, 4.7E+04) | (0.6, 2.9) | (1.1E+06, 1.3E+06) |
| Variance explained (%) | 32.4 | 1.0 | 0.0 | 66.6 |
| **gpiCor** |  |  |  |  |
| Estimate | 3.7E+05 | 3.0E+04 | 2.5 | 1.0E+06 |
| HPDI | (2.7E+05, 4.8E+05) | (0.0, 6.4E+04) | (0.9, 4.6) | (9.6E+05, 1.1E+06) |
| Variance explained (%) | 26.1 | 2.1 | 0.0 | 71.8 |
| **iSize** |  |  |  |  |
| Estimate | 1.0E+07 | 3.1E+06 | 1.5 | 5.8E+07 |
| HPDI | (7.0E+06, 1.4E+07) | (0.0, 5.7E+06) | (0.6, 2.9) | (5.4E+07, 6.2E+07) |
| Variance explained (%) | 14.6 | 4.3 | 0.0 | 81.1 |

Note: HPDI, 95% Highest Posterior Density Interval.

**Supplementary table 2.** Results from multiple linear regression analyses

| model | *p* value |  |  |  |
| --- | --- | --- | --- | --- |
|  | iSize<br>GEBVs | gpi<br>GEBVs | gpiCor<br>GEBVs | gpdCor<br>GEBVs |
| gpd = iSize GEBVs + gpi GEBVs + e | *** | N.S. | - | - |
| gpd = gpi GEBVs + e | - | ** | - | - |
| gpd = iSize GEBVs + gpiCor GEBVs + e | *** | - | N.S. | - |
| gpd = gpiCor GEBVs + e | - | - | ** | - |
| gpd = iSize GEBVs + gpdCor GEBVs + e | *** | - | - | N.S. |
| gpd = gpdCor GEBVs + e | - | - | - | N.S. |

N.S., Not significant; \*, $p < 0.05$ ; \*\*, $p < 0.01$ ; \*\*\*, $p < 0.001$

**Supplementary table 3.** GWAS candidate genes. SNP position interval denotes the most significant SNPs in the peak coloured in **Figure 5A-D**. %identity refers to the identity with the closest *Medicago truncatula* homolog. The annotation column shows the species the gene was annotated based on; [Mt]: *Medicago truncatula*, [Tp]: *Trifolium pratense*, [Ca]: *Cicer arietinum*.

| Chr. | SNP position | Gene | Gene position |  | Annotation | %identity | Phenotypic trait(s) |
| --- | --- | --- | --- | --- | --- | --- | --- |
|  |  |  | Start | End |  |  |  |
| 1 | 1910067 - 1909821 | chr1.jg240.t1 | 1909818 | 1918815 | small RNA degrading nuclease [Mt] | 86.36% | iSize |
| 3 | 28987454 - 28989233 | 51 bp from stop codon: chr3.jg4449.t1 | 28989826 | 28987505 | u6 snRNA-associated-like-Smprotein [Mt] | 73.83% | iSize+gpd |
| 6 | 27501506 | chr6.jg4045.t1 | 27504540 | 27500993 | E3 ubiquitin-protein ligase rhf2a-like protein [Tp] | 86.89% | iSize |
| 7 | 19892133 - 20181795 | chr7.jg3208.t1 | 20181419 | 20182720 | extracellular dermal glycoprotein [Mt] | 88.45% | gpd |
| 7 | 29979844 - 30231861 | 342bp from stop codon: chr7.jg4843.t1 | 30236357 | 30232200 | bag family molecular chaperone regulator 4-like protein [Tp] | 80.09% | gpd |
| 7 | 51953440 - 51953748 | upstream 790bp: chr7.jg8266.t1 | 51958855 | 51955400 | WD repeat-containing protein GTS1 [Ca] | 86.89% | iSize+gpd |
| 8 | 32983608 - 32988584 | chr8.jg4570.t1 | 32989033 | 32984274 | kinetochore protein spc25-like [Tp] | 92.41% | gpd |
| 13 | 4876408-4876434 | 66bp from stop codon: chr13.jg754.t1 | 4879246 | 4876474 | transcription factor S-II, central domain protein [Mt] | 83.47% | iSize |

**Supplementary table 4.** Composition of peat used for the greenhouse experiments.

| Ingredient | Quantity added per m<sup>3</sup> |
| --- | --- |
| Sphagnum | 0.8 m<sup>3</sup> |
| N-P-K-0-7-22 + micro | 1.2 kg |
| Superphosphate (crushed) | 0.4 kg |
| Micromax | 0.05 kg |
| Calcium | 2.4 kg |
| Perlite type 3 (2-6 mm) | 200 l |
| BARA clay (2-6 mm) | 60 kg |
| NO<sub>3</sub>-N | 0.00 g |
| K | 268.80 g |
| Mo | 3.27 g |
| Zn | 1.10 g |
| NH<sub>4</sub>-N | 0.00 g |
| Mg | 59.10 g |
| Cu | 2.42 g |
| Fe | 8.94 g |
| P | 117.60 g |
| B | 0.58 g |
| Mn | 3.77 g |
| S | 59.00 g |

**Supplementary table 5.**
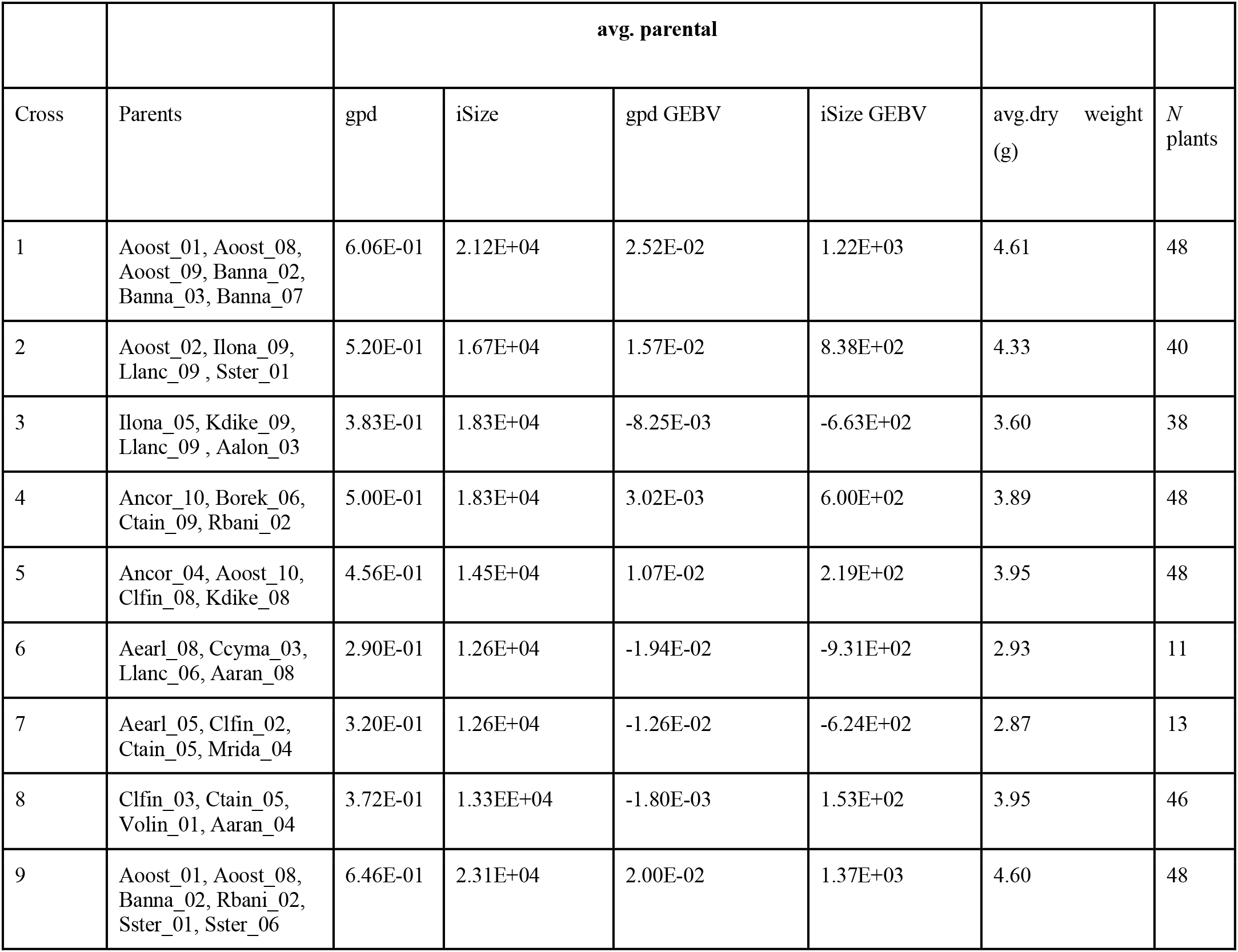
Overview of F1 populations with information about their average dry weights, number of established plants, parents used to generate the polycrosses and their average iSize, gpd, GEBV.

|  |  | avg. parental |  |  |  |  |  |
| --- | --- | --- | --- | --- | --- | --- | --- |
| Cross | Parents | gpd | iSize | gpd GEBV | iSize GEBV | avg.dry weight<br>(g) | N<br>plants |
| 1 | Aoost_01, Aoost_08,<br>Aoost_09, Banna_02,<br>Banna_03, Banna_07 | 6.06E-01 | 2.12E+04 | 2.52E-02 | 1.22E+03 | 4.61 | 48 |
| 2 | Aoost_02, Ilona_09,<br>Llanc_09, Sster_01 | 5.20E-01 | 1.67E+04 | 1.57E-02 | 8.38E+02 | 4.33 | 40 |
| 3 | Ilona_05, Kdike_09,<br>Llanc_09, Aalon_03 | 3.83E-01 | 1.83E+04 | -8.25E-03 | -6.63E+02 | 3.60 | 38 |
| 4 | Ancor_10, Borek_06,<br>Ctain_09, Rbani_02 | 5.00E-01 | 1.83E+04 | 3.02E-03 | 6.00E+02 | 3.89 | 48 |
| 5 | Ancor_04, Aoost_10,<br>Clfin_08, Kdike_08 | 4.56E-01 | 1.45E+04 | 1.07E-02 | 2.19E+02 | 3.95 | 48 |
| 6 | Aearl_08, Ccyma_03,<br>Llanc_06, Aaran_08 | 2.90E-01 | 1.26E+04 | -1.94E-02 | -9.31E+02 | 2.93 | 11 |
| 7 | Aearl_05, Clfin_02,<br>Ctain_05, Mrida_04 | 3.20E-01 | 1.26E+04 | -1.26E-02 | -6.24E+02 | 2.87 | 13 |
| 8 | Clfin_03, Ctain_05,<br>Volin_01, Aaran_04 | 3.72E-01 | 1.33EE+04 | -1.80E-03 | 1.53E+02 | 3.95 | 46 |
| 9 | Aoost_01, Aoost_08,<br>Banna_02, Rbani_02,<br>Sster_01, Sster_06 | 6.46E-01 | 2.31E+04 | 2.00E-02 | 1.37E+03 | 4.60 | 48 |

## Supplementary Figures

**Supplementary figure 1.**
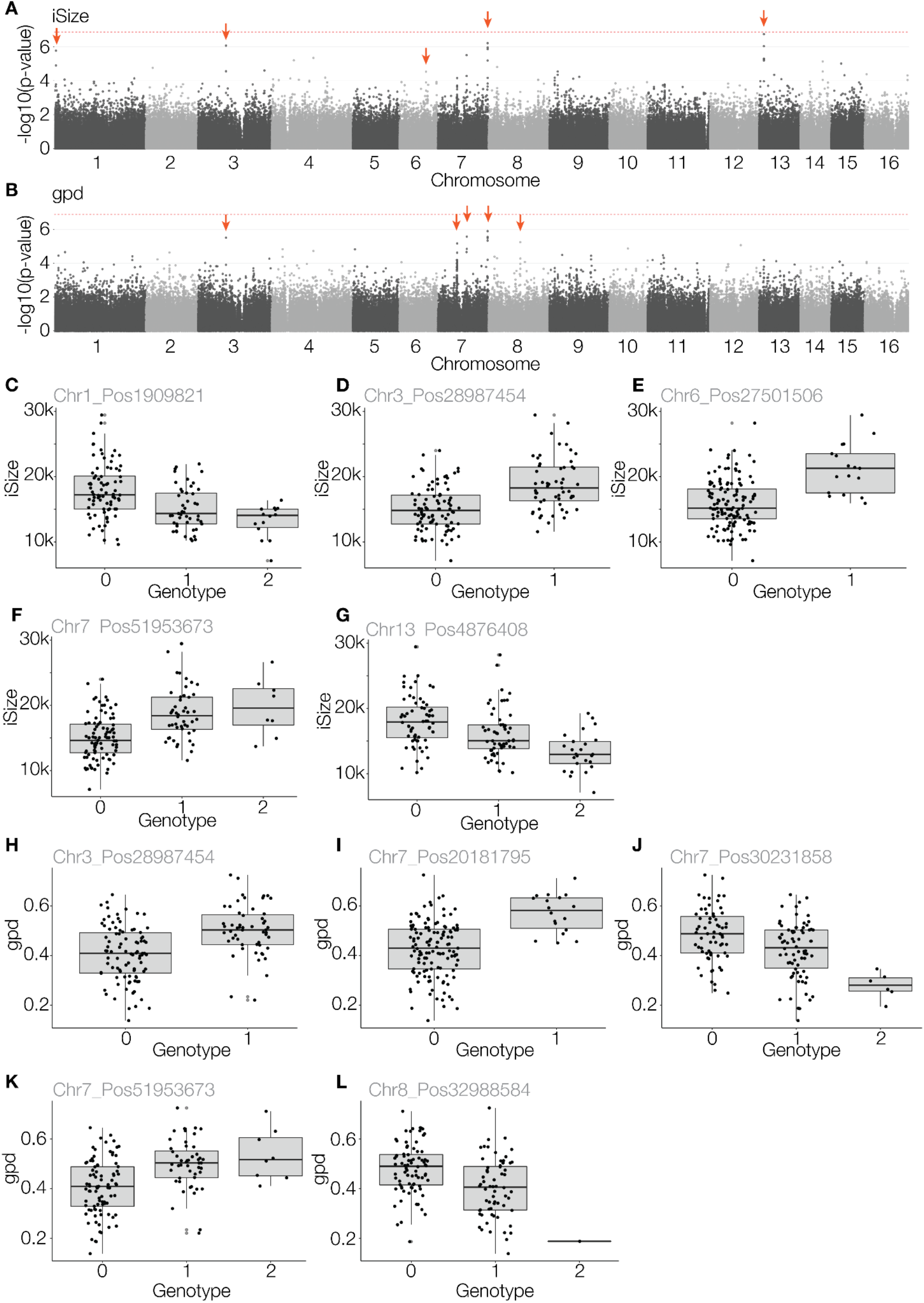
**A-B:** Manhattan plots showing results of GWAS of **A:** iSize or **B:** gpd (*n* = 145). The genetic model is set to diploid. The red dotted line indicates the Bonferroni threshold at 6.9. Effect plots for the most significant SNP for each peak indicated with an orange arrow is shown in **C-L**. **C-G:** Effect plots for the most significant SNP in each peak for iSize. **H-L:** Effect plots for the most significant SNP in each peak for gpd.

**Supplementary figure 2.**
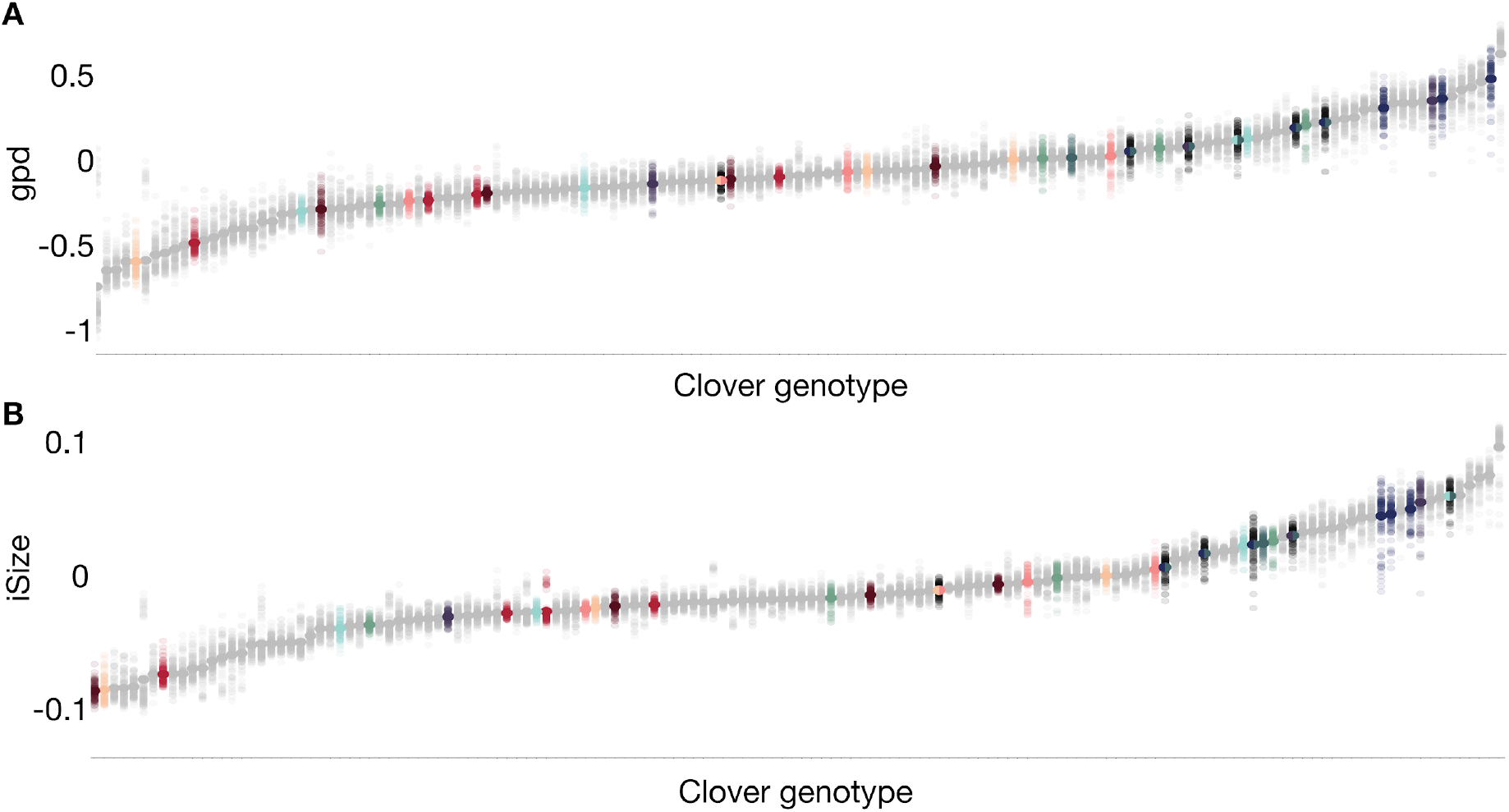
F1 parental distribution of GEBVs of gpd (**A**) or iSize (**B**). Colors refer to Figure 6.

**Supplementary figure 3.**
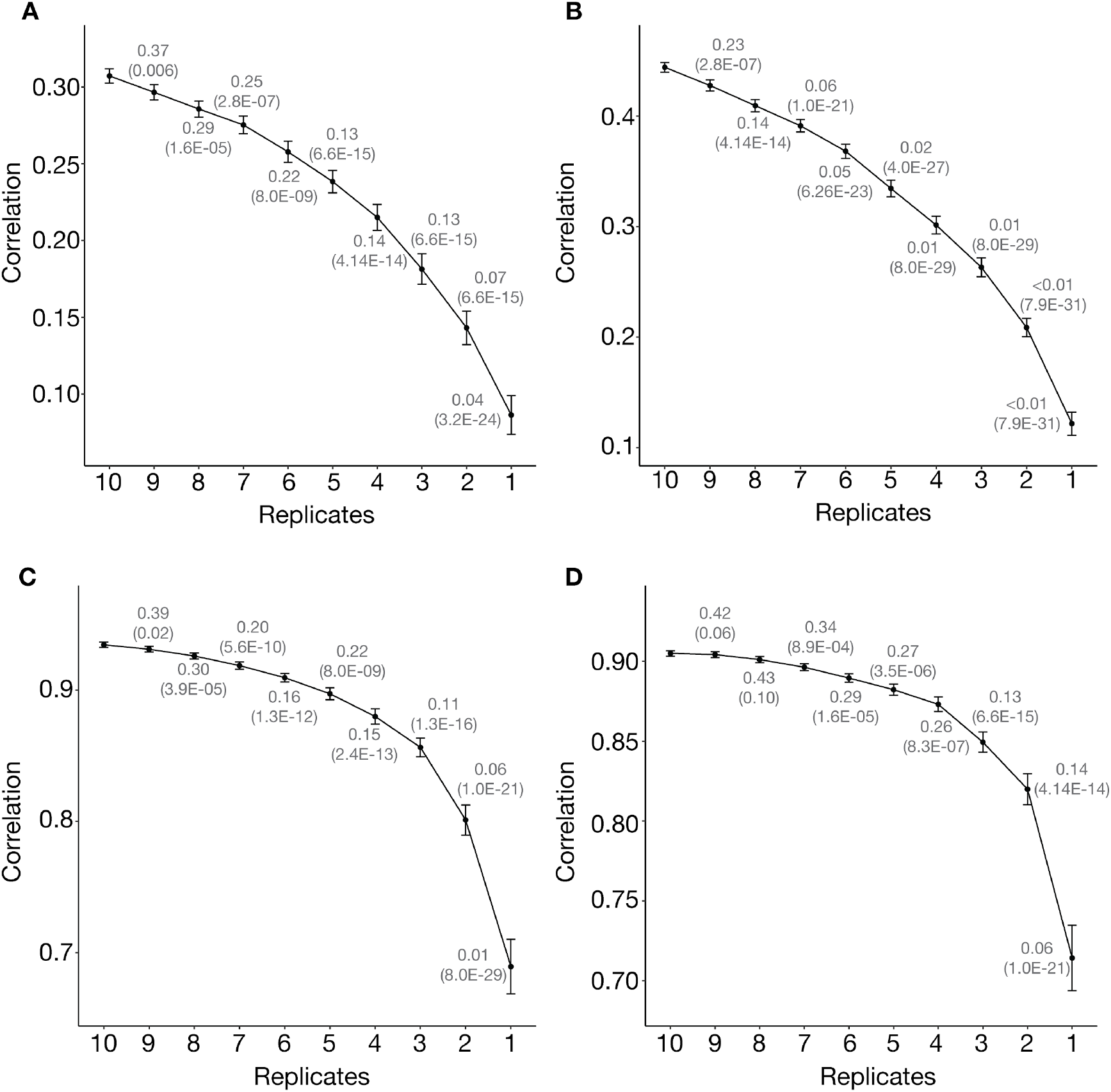
Replicate reduction. Correlation between GEBVs and phenotypes for the dataset using only clover genotypes with at least 10 replicates. Error bars display standard errors. The fractions above the data points refer to the frequency that using the indicated number of replicates performed better than using 10 replicates for prediction. The numbers in parentheses indicate the *p* value that the indicated number of replicates led to a prediction performance equal to that of using 10 replicates (paired sample sign test). **A-B:** F0 prediction of gpd using (**A**) gpd and (**B**) iSize. **C-D:** F1 prediction of gpd using (**C**) gpd and (**D**) iSize

**Supplementary figure 4.**
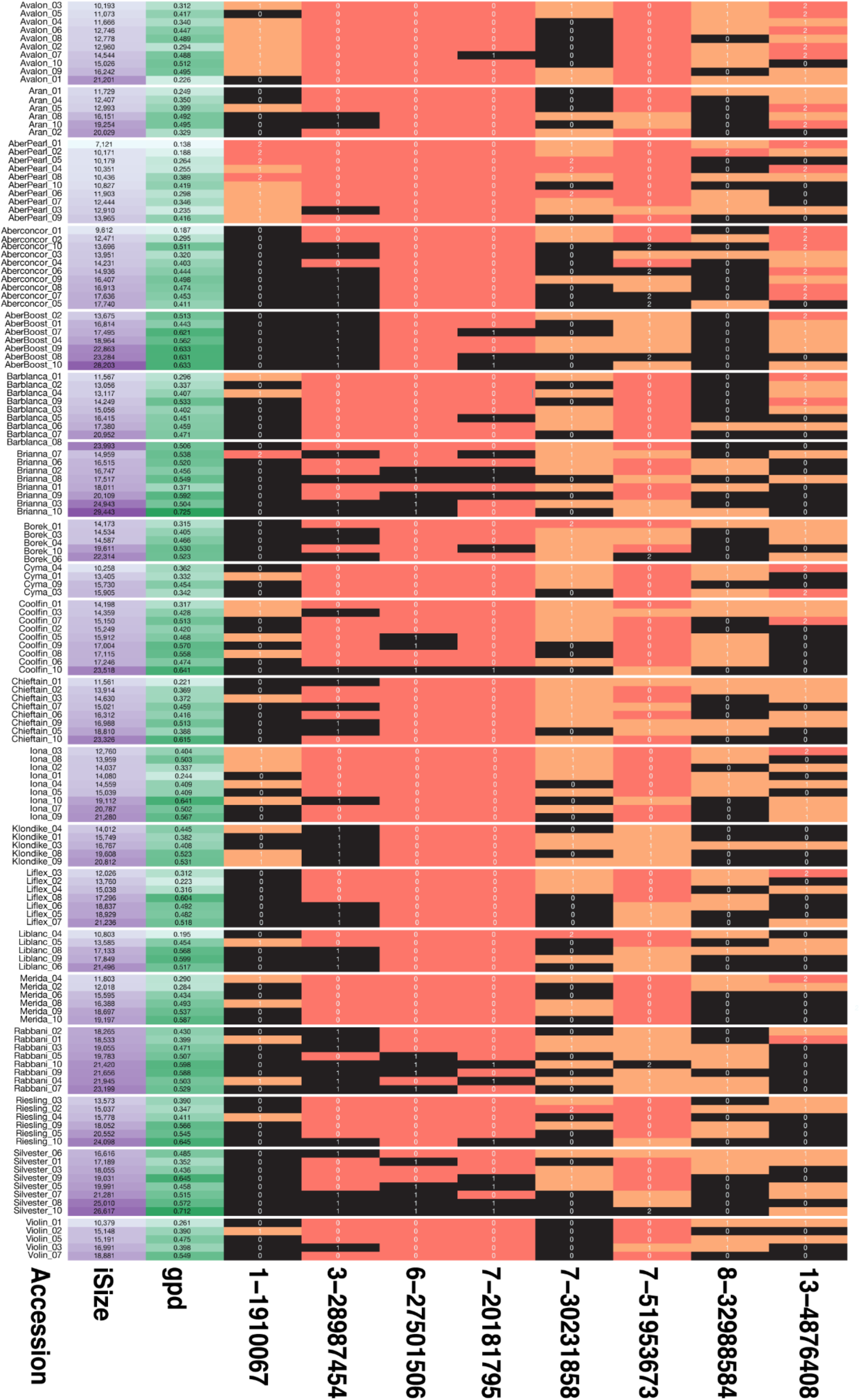
Variation in QTL genotypes (0, homozygous reference; 1, heterozygote; 2, homozygous alternative. Each line reports the genotype, average iSize, gpd and the clover genotype. Genotypes are coloured by their effect on iSize. Black blocks indicate the allele with the largest median iSize, red blocks indicate the allele with the lowest median iSize, orange blocks indicate heterozygosity. Accessions are grouped into blocks row-wise according to variety and sorted within variety according to iSize.

## Supplementary files

**Supplementary file 1:** Raw growth data

**Supplementary file 2:** Observations of single plants data

**Supplementary file 3:** Average phenotypes and GEBVs

**Supplementary file 4:** Imputed genotype file

**Supplementary file 5:** GWAS results, top 200 most significant SNPs

**Supplementary file 6:** Dry weight of F1 plants

